# In situ architecture of the lipid transport protein VPS13C at ER-lysosomes membrane contacts

**DOI:** 10.1101/2022.03.08.482579

**Authors:** Shujun Cai, Yumei Wu, Andres Guillen-Samander, William Hancock-Cerutti, Jun Liu, Pietro De Camilli

## Abstract

VPS13 is a eukaryotic lipid transport protein localized at membrane contact sites. Previous studies suggested that it may transfer lipids between adjacent bilayers by a bridge-like mechanism. Direct evidence for this hypothesis from a full-length structure and from EM studies *in situ*, however, is still missing. Here we have capitalized on AlphaFold predictions to complement the structural information already available about VPS13 and to generate a full-length model of human VPS13C, the Parkinson’s disease-linked VPS13 paralog localized at contacts between the ER and endo/lysosomes. Such model predicts a ~30-nm rod with a hydrophobic groove that extends throughout its length. We further investigated whether such a structure can be observed *in situ* at ER-endo/lysosome contacts. To this aim, we combined genetic approaches with cryo-focused-ion-beam (cryo-FIB) milling and cryo-electron tomography (cryo-ET) to examine HeLa cells overexpressing this protein (either full length or with an internal truncation) along with VAP, its anchoring binding partner at the ER. Using these methods we identified rod-like densities that span the space separating the two adjacent membranes and that match the predicted structures of either full length VPS13C or its shorter truncated mutant, thus providing the first *in-situ* evidence for a bridge-model of VPS13 in lipid transport. Intriguingly, the majority of the VPS13C rods were separated from the ER membranes by a narrow gap, suggesting that while VAP anchors the protein to the ER, direct contact of the VPS13C rod with the ER bilayer to allow lipid transport may be independently regulated.

## Introduction

Communication between subcellular membranous organelles mediated by direct contacts not leading to their fusion plays a critical role in the physiology of eukaryotic cells and is an important complement to interorganelle communication mediated by vesicular transport. The endoplasmic reticulum (ER), in particular, establishes direct contacts with all other membranous organelles of the cell^1^. One of the functions of such contacts is to mediate lipid transfer between the two closely apposed membranes via proteins that contain lipid transport modules and also act as direct or indirect tethers between them^2–4^. Until few years ago, lipid transfer proteins acting at membrane contact sites in eukaryotic cells were thought to act exclusively by a shuttle mechanism, i.e. via modules that harbor one or few lipids and shuttle back and forth between the two participating membranes^5^. However, recent studies have also suggested the occurrence of proteins that provide hydrophobic bridges along which lipids can move directly from one bilayer to another to achieve bulk lipid transport^5, 6^.

The founding member of this class of protein is VPS13, a protein first identified in yeast by genetic screens for membrane traffic proteins^7–9^. Subsequently, VPS13 was putatively linked to lipid transport because of the identification of VPS13 dominant mutations as suppressors of the deficiency of ERMES (ER–Mitochondria Encounter Structure), a protein complex that mediates lipid transport between the ER and mitochondria in yeast^10, 11^. The human genome comprises four VPS13 proteins, referred to as VPS13A, VPS13B, VPS13C and VPS13D, which have different localizations at membrane contact sites and whose mutations result in neurodevelopmental defects or neurodegenerative conditions^12–15^. A role of VPS13 family proteins in lipid transport, and more specifically at membrane contact sites, has now been supported by genetic, biochemical, imaging and structural studies^6, 10, 11, 16–21^. Fragments of VPS13 were shown to contain multiple lipids and to transfer lipids between artificial liposomes^17^. Crystallographic studies of N-terminal region of VPS13 of Chaetomium thermophilum revealed a hydrophobic cavity^17^ which cryo-EM studies of a longer rod-like fragment of the same protein subsequently suggested to continue as a hydrophobic groove along the entire length of the rod^19^. Moreover, binding sites for the ER and for other organelles have been identified in the N-terminal and C-terminal region of several VPS13 family members, respectively^17, 18, 20–22^. Together, these results have suggested a bridge model of lipid transport ideally suited for the bulk delivery of lipids and in line with some of the proposed functions of VPS13 family members in membrane expansion^6^. The identification of regions of amino acid similarity and of predicted structural similarities between VPS13 and the autophagy factor ATG2 had suggested a similar function of ATG2^17, 23, 24^.

Despite these studies, direct evidence for an arrangement of VPS13-ATG2 family proteins *in situ* that would support a bridge model of lipid transport is still missing. Goal of this study was to address this gap by combining cryo-focused-ion-beam (cryo-FIB) milling and cryo-electron tomography (cryo-ET) imaging, the a recently developed methodology that allows to resolve the in situ high-resolution (sub-nanometer) structure of proteins of interest^25–27^. To this aim we focused on VPS13C, the VPS13 isoform that is localized at ER-late endosome/lysosomes (endo/lysosomes) contact sites via its FFAT motif-dependent interaction with the ER protein VAP and with Rab7 on endo/lysosomal membranes. Overexpression of VPS13C and VAP results in the extensive formation or ER-endo/lysosome contacts, thus facilitating their identification in cryo-electron tomograms. Our cryo-ET analysis of cell cryo-lamellae generated by cryo-FIB milling provides the first *in situ* structural evidence to support the bridge model of lipid transport and also suggests a dynamic interaction of VPS13C with the ER bilayer. VPS13C loss-of-function results in lysosomal dysfunction at the cellular level^21^ and causes familial Parkinson’s disease in human patients^14^. Moreover, as dysfunction of other VPS13 family also results in neurological diseases^28^, a better understanding of in situ architecture VPS13C has implications for both biology and medicine.

## RESULTS

### Structure of VPS13C as predicted by AlphaFold

As a premise to the identification of VPS13C by cryo-ET, we took advantage of AlphaFold v2.0 to gain insight into its full-length atomic structure^29^. Human VPS13C isoform 1 comprises 3753 amino acids (a.a.). Since predictions of amino acid sequences longer that 2000 a.a. by current AlphaFold algorithms are inaccurate, we predicted the structure of three segments of the protein, each one containing ~600 a.a. overlap with the adjacent segment, and then aligned them to generate a full-length structure (Fig. 1B, Movie S1). Such structure is represented by a 29.3 nm long rod whose backbone is a narrow twisted β-sheet running along its entire length. The β-sheet, which is flanked by α-helices and disordered loops forms the floor of a hydrophobic groove that extends throughout the rod (Fig. 1C-D, Movie S2) and thus could mediate the sliding of lipids from one end to the other end of the protein. The presence of this groove is clearly visible in a view perpendicular to the axis of the rod (Fig. 1C) where the groove appears as a tubular cavity given the twisting of the rod. One of the disordered loops surrounding the rod contains the FFAT consensus for binding to VAP and, accordingly, the AlphaFold-Multimer algorithm^30^ maps to this loops such interaction (Fig. 1B, Movie S1). A region of VPS13 downstream to the so-called VAB domain (Fig. 1A), which had previously been described as the APT-1 domain (Fig. 1A), comprises the C-terminal portion of the twisted long β-sheet and is directly continuous with the portion of this sheet preceding the VAB domain, so that VAB domain represents an outpocketing of the rod (Fig. 1B, 1E). The β-sheet portion of the APT-1 domain is in turn interrupted by a smaller outpocketing of about 67 a.a. which has sequence and fold similarities to WWE domains (Figs. 1F, S1). This small domain, not previously described in VPS13, is also present in VPS13A (but not VPS13B and VPS13D) and in several VPS13 proteins across evolution^31^. The AlphaFold predicted shape of the N-terminal approximately 330 a.a. of the protein, which comprise the so-called chorein_N motif (~a.a. 1-120), is in good agreement with the fold of this region from Chaetomium thermophilum as determined by crystallography, although it is somewhat wider^17^. Likewise, the AlphaFold model generally agrees with the cryo-EM structure of the first 1390 a.a. of VPS13 from Chaetomium thermophilum, although with a tighter twisting. The VAB domain, which contains 6 repeats of a module entirely composed of β-sheets^18^, forms an arc structure which likely corresponds to the loop structure previously observed in a low resolution negative staining EM images of yeast VPS13^16^ (Fig. 1G). In the AlphaFold the predicted structure the arc is arranged in a plane roughly perpendicular to the axis of rod, while in the De et al study several orientations of arc relative to the rod were reported. Thus, a flexible connection of this region relative to the rod with the possibility of large-scale rotations had been suggested^16^, as now confirmed by the AlphaFold full-length structure. Finally, AlphaFold confirms a C-terminal PH domain preceded by a bundle of four alpha helices similar to the C-terminal region of ATG2 (ATG-C), a region that we had previously called as DH-L because of a distant similarity to DH domains (Fig. 1H). Having defined the predicted structure of VPS13C, we set out to build on this information to explore the presence of structures that fit these predictions and their arrangement at membrane contact sites in intact cells.

**Figure 1.**
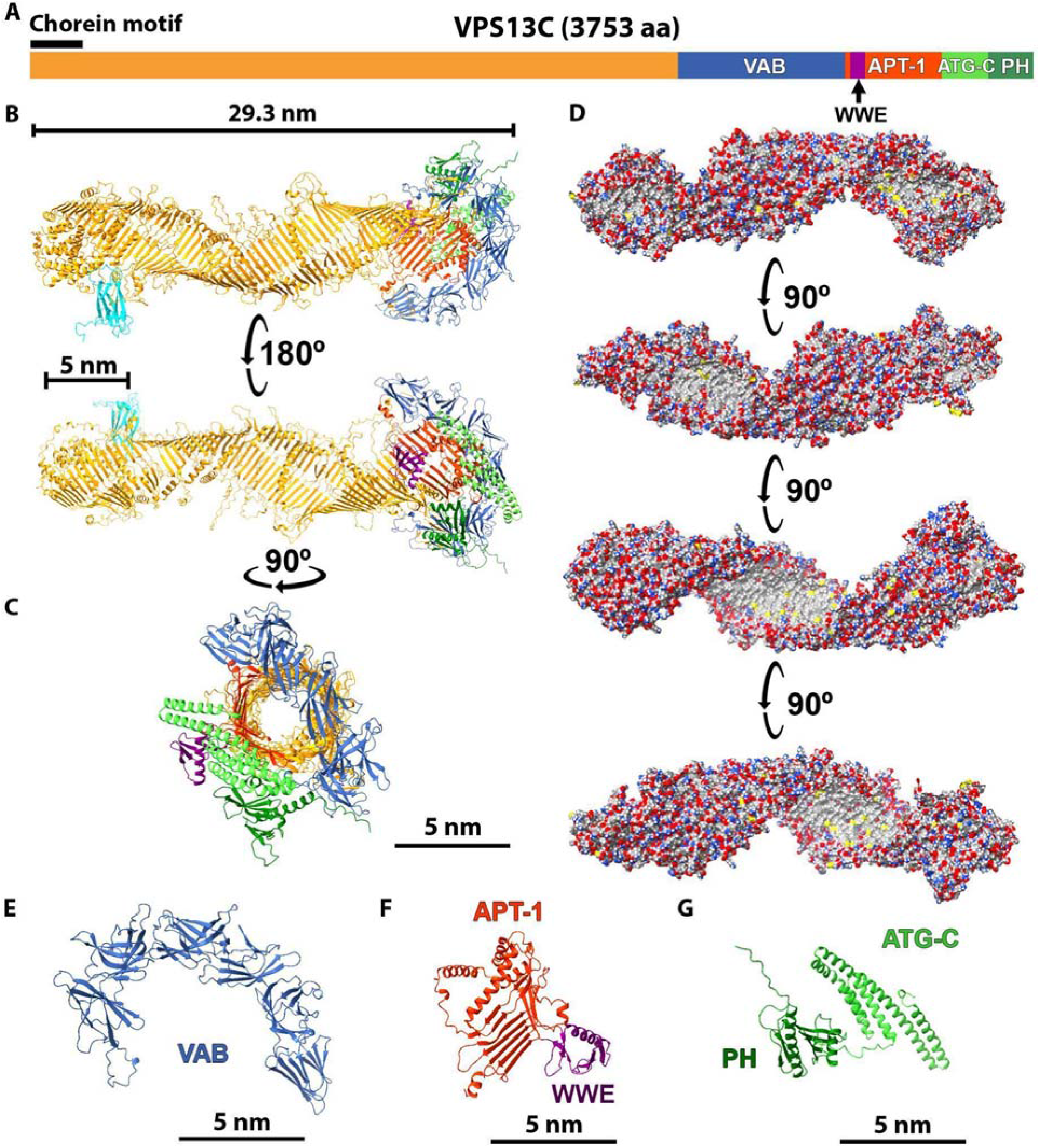
Structure of VPS13C and its binding partners as predicted from AlphaFold. (A) Schematic cartoon of the domain architecture of human VPS13C. (B and C) Predicted structure of fulllength VPS13C in two perpendicular views. The color scheme is consistent with the one shown in panel A. The MSP domain of VAPA bound to VPS13C is also shown (in cyan) in field B. The grove that runs along the entire protein appears as a tunnel due to the twisting of the elongated β-sheet (D) Surface representations of the protein (90 degree rotation interval) showing carbon atoms in grey (hydrophobic surfaces), oxygens in red (negative charges), nitrogens in blue (positive charges) and sulfur in yellow. Note the presence of a continuous hydrophobic groove (grey) along the protein. For clarity of presentation, some α-helices and disordered loops are not shown. (E-G) Enlarged views of the individual C-terminal domains of the protein. The APT-1 domain forms the C-terminal region of the elongated β-sheet that represents the core of the protein.

### A cellular model for *in-situ* analysis of VPS13C-mediated contacts

To locate VPS13C-mediated membrane contacts for high resolution cryo-ET imaging, we first developed a cellular system enriched in such contacts. Overexpression of VPS13C^Halo and GFP-VAP (VAP-B) with or without Rab7 (possibly because Rab7 is not present at limiting concentration) is sufficient to produce a massive expansion of the normally occurring ER to endo/lysosome contacts (as represented schematically in Fig. 2A), resulting in endo/lysosomes completely surrounded by ER cisterns, as shown by fluorescence microscopy^17, 21^ (Fig. 2B) and electron microscopy (Fig. 2C). Frequently this ER is “thin” ER. i.e. ER virtually devoid of a lumen so that the two opposite membranes of the cisterns are in tight apposition to each other, as often observed at contact sites of the ER with other membranes^32, 33^. Endo/lysosomes completely surrounded by ER can also be observed upon overexpression of other proteins that act as tethers between these two organelles^34, 35^. One of them is PDZD8, which has a N-terminal transmembrane region anchored in the ER, binds Rab7 on endo/lysosomes and transports lipids via an SMP domain. Interestingly, co-expression of VPS13C and PDZD8 show that the two proteins segregate to distinct domains within the contact area between the two organelles (Fig. 2D). Furthermore, correlative fluorescence microscopy and focused ion beam-scanning electron microscopy (FIB-SEM) not only confirmed that VPS13C and PDZD8 enriched sites corresponded to contacts between endo/lysosomes and the ER, but also revealed a striking difference in the distance between the two organelles at the two domains (Fig. 2E, 2E’, 2F, Movie S3). Although the resolution of FIB-SEM did not allow a precise measurement of such distances, the intermembrane space at VPS13C positive domains was much greater than the distance observed at PDZD8-dependent ER to endo/lysosome contacts^34^ (Fig. 2E, 2E’, 2F). The concentration of overexpressed VPS13C at extended ER-endo/lysosome contacts suggested that VPS13C and VAP overexpressing cells represent an optimal cellular system to analyze VPS13C-mediated membrane contacts at high resolution by a combination of cryo-FIB milling and cryo-ET imaging.

**Figure 2.**
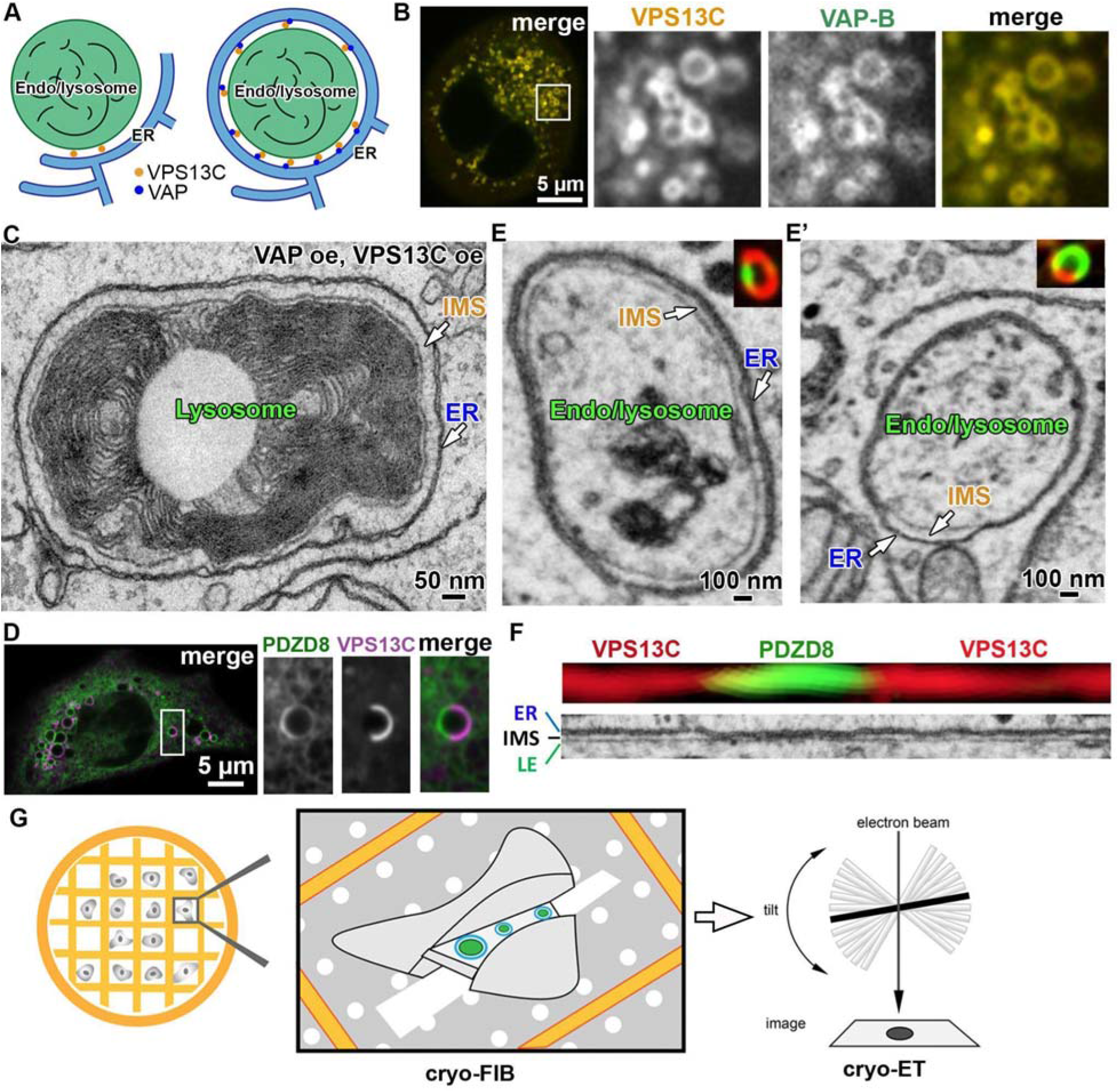
Experimental system used to visualize VPS13C *in situ* with cryoET. (A) Cartoons showing VPS13C-mediated ER-endo/lysosome contacts in wildtype cells (left) and in cell over-expressing VPS13C and VAP-B (right). (B) Left: “merged” confocal microscopy z-slice image of a HeLa cell over-expressing VPS13C^Halo and GFP-VAP. The region enclosed by a box is shown at fivefold enlargement in the split and merged channels at right. The majority of VPS13C^Halo positive endo/lysosomes are also positive for the ER protein VAP, demonstrating enwrapping by ER. (C) Electron micrograph of a lysosome completely surrounded by ER (thin ER) from a COS-7 cell overexpressing VPS13C and VAP-B. IMS: the inner-membrane space between the ER and the lysosome membrane. (D) Left: “merged” confocal microscopy z-slice images of a Cos-7 cell over-expressing VPS13C^Halo and PDZD8-EGFP. The region enclosed by a box is shown at higher magnification in the split and merged channels at right. (E and E’) Correlative fluorescence (insets) and FIB-SEM images (large fields) of endo/lysosomes from Cos-7 cells overexpressing VPS13C^halo (red) and PDZD8-EGFP (green). VPS13C (red) and PDZD8 (green) segregate to distinct domains within the contact area between ER (thin ER) and endo/lysosomes. IMS: the inner-membrane space between the ER and the lysosome membrane. The space between the two organelles is much wider at the contacts where VPS13C is localized. (F) Linearized membrane contacts from the same endo/lysosome of panel E (processed with Fiji Kymograph) showing the difference width of the intermembrane space (IMS) at VPS13C- and PDZD8-positive sites. (G) Cryo-electron tomography workflow. Cells seeded on EM grids were first examined by live confocal microscopy to locate cells overexpressing VPS13C and VAP. Then grids were plunge frozen to maintain the life-like state of cells. ~150-nm-thick cryo-lamellae were generated by cryo-FIB and imaged with cryo-ET.

### VPS13C-mediated membrane contacts as visualized with cryo-ET

A schematic cartoon of the flow of work to analyze membrane contact sites mediated by VPS13C is shown in Fig. 2G. Cells were grown on a gold-mesh EM grid, cotransfected with VPS13C^Halo and GFP-VAP-B, and examined by live fluorescence microscopy to identify a cell co-expressing both proteins, where vesicular structures surrounded by both Halo and GFP fluorescence (expected to represent expanded ER-endo/lysosome contacts) could be observed (Fig. 2B). The grid was then plunge-frozen and the selected cell was processed by cryo-FIB milling to generate an approximately 150 nm-thick cryo-lamellae to be analyzed by Cryo-ET (Fig. 2G). As a negative control, cryo-ET was performed on cells expressing VPS13C^Halo alone, GFP-VAP alone or neither protein (wild type cells).

Abundant presence of endo/lysosomes nearly completely surrounded by ER were observed in cryo-lamellae of cells co-overexpressing VPS13C and VAP-B (Fig. 3A and A’) but not in the cryo-lamellae of control cells (Fig. 3B-D, 3F-H). At the massive contacts observed in VPS13C and VAP-B expressing cells, the distance between the two membranes was relatively uniform (Fig. 3A, 3E). A violin plot of membrane spacing measured at multiple sites around different endo/lysosomes showing a peak at ~33.5 nm (Fig. 4C). The same massive ER wrapping (with a similar width of the intermembrane space) was also occasionally observed around organelles enclosed by a double membrane raising the possibility that they may represent amphisomes (Fig. S2A).

**Figure 3.**
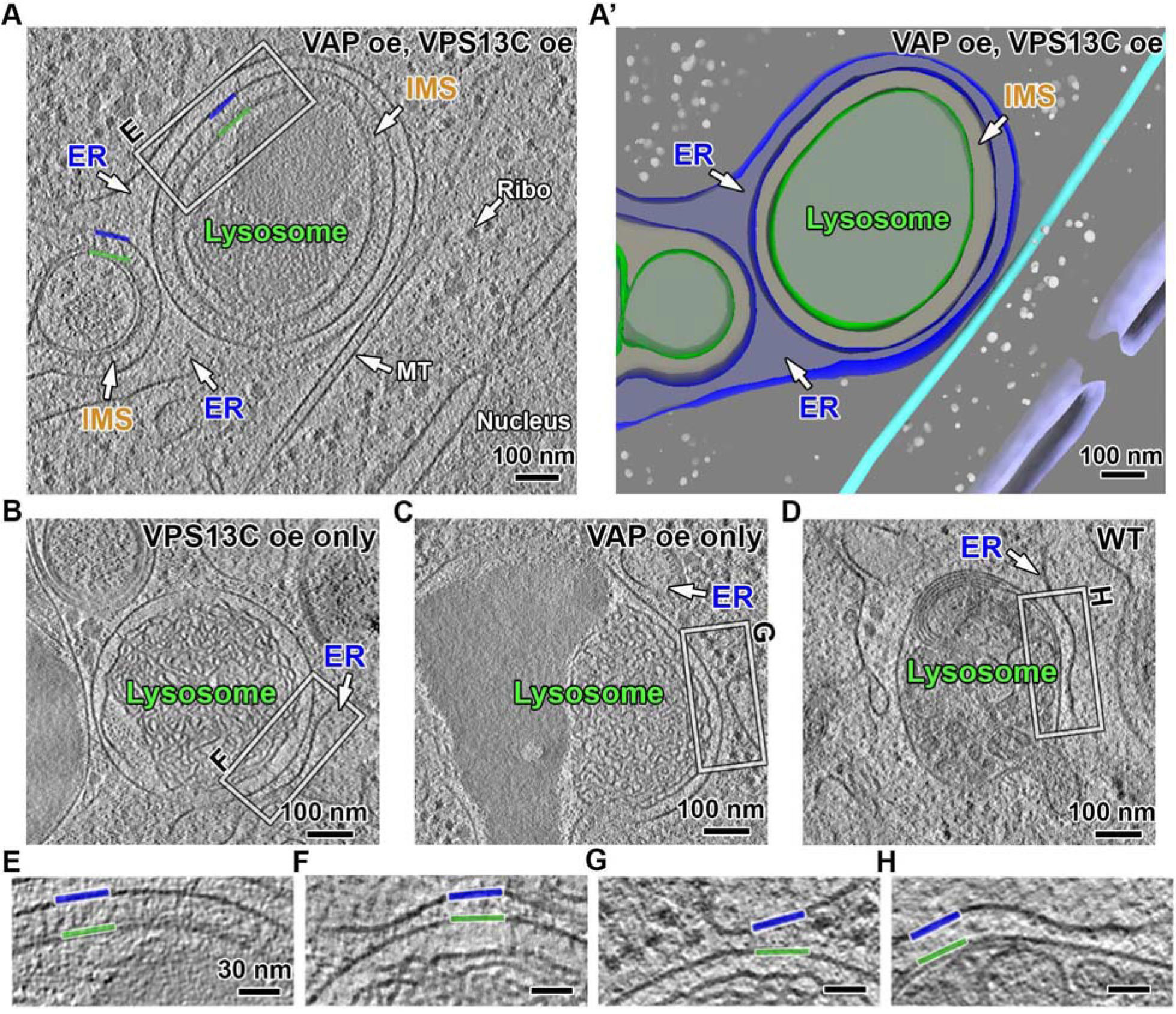
VPS13C-mediated membrane contact sites. (A) Cryo-tomographic slice (1-nm thick) showing VPS13C mediated membrane contacts between lysosomes and ER in a VPS13C and VAP-B co-overexpressing HeLa cell. IMS: inner-membrane space between the ER and the lysosome membrane. (A’) 3D view of the tomogram shown in panel A. Grey sphere: ribosome, cyan tube: mitochondria, purple: nuclear envelope (B-D) Negative controls showing contacts between lysosomes and ER in cells without VPS13C and VAP co-overexpression. (E-H) Enlarged views (2 fold) of the region boxed in panels A-D showing ER-lysosome contacts. Blue and green lines: ER and lysosome membrane, respectively.

**Figure 4.**
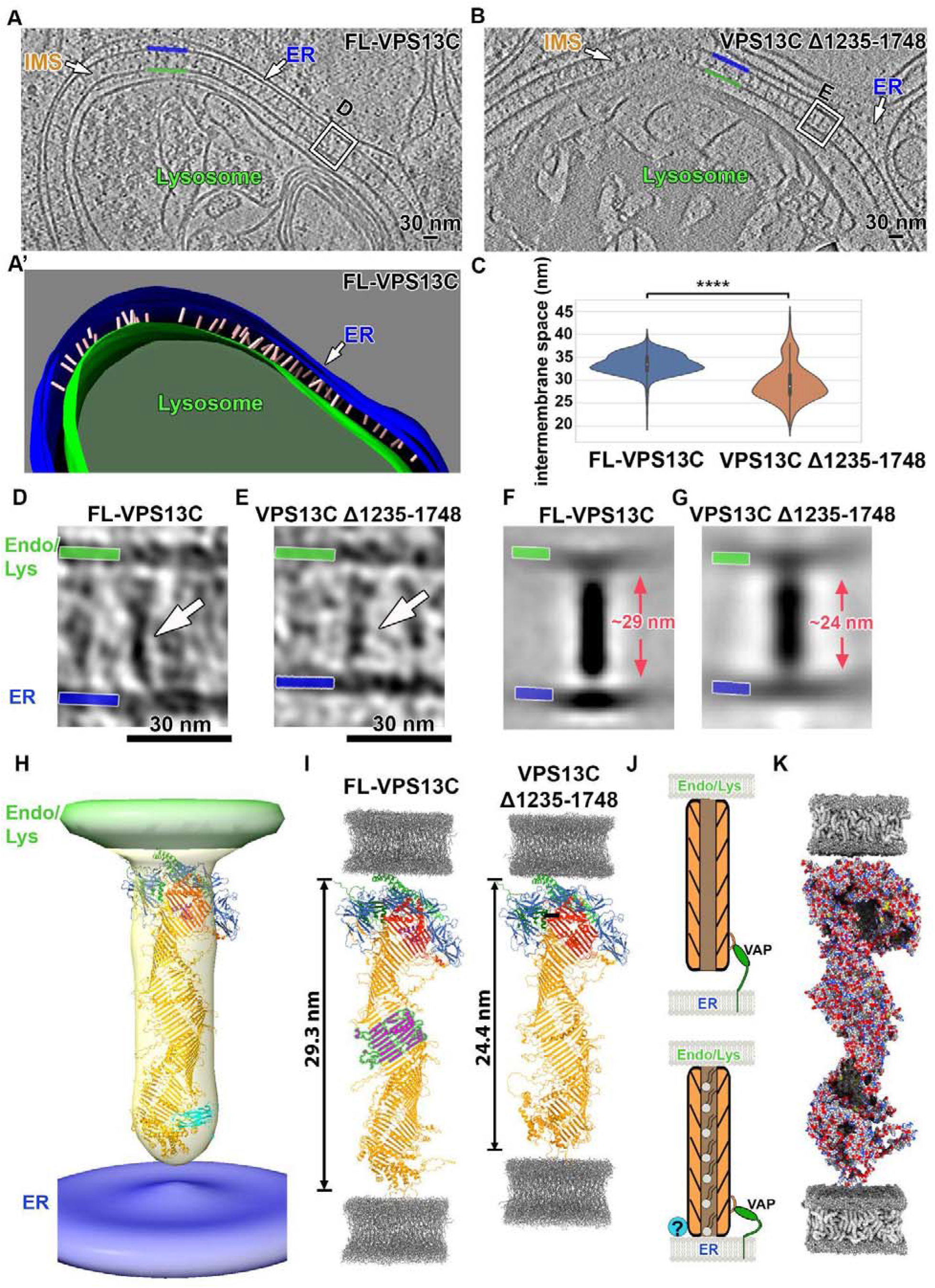
Rod-like structures with the expected length of VPS13C bridge the membranes at contacts between the ER and lysosomes. (A) Cryotomographic slice (1-nm thick) showing abundant rod-shaped densities at ER-lysosome contacts in a HeLa cell overexpressing full length VPS13C and VAP. Blue and green lines: ER and lysosome membrane, respectively. IMS: intermembrane space between the ER and the lysosome membrane. (A’) 3D segmentation view of panel A. Pink: VPS13C-like rod densities. (B) Cryotomographic slice (1-nm thick) showing shorter rod-shaped densities and narrower intermembrane space relative to field A, at ER-lysosome contacts in a HeLa cell overexpressing a truncated VPS13C mutant (VPS13C Δ1235-1748) and VAP. (C) Violin plot showing the distribution of ER-lysosome intermembrane distances at contacts mediated by full-length VPS13C and VPS13C Δ1235-1748, respectively. **** p<0.0001. (D and E) Enlarged views of the regions boxed in panel A (in a different tomographic slice) and panel B, respectively, showing rod-shaped densities bridging the ER membrane (blue) to the lysosome membrane (green). (F and G) Subtomogram-average density maps showing a 29-nm-long full length VPS13C rod (panel F) and a 24-nm-long VPS13C Δ1235-1748 rod (panel G) bridging the two adjacent membranes. C100 symmetry was applied to enhance signal to noise ratio. See Fig. S4A for averages without applying symmetry. (H) Three-dimensional view of panel F. The density corresponding to VPS13C is shown in light orange, fitted with the full-length VPS13C predicted structure from AlphaFold v2.0. The color scheme of predicted structure is consistent with the one shown in Fig. 1A. (I) Predicted structures of full-length VPS13C (left) and VPS13C truncation mutant (Δ1235-1748) in between two lipid bilayers. The fragment shown in magenta is removed from full-length VPS13C to generate the truncation mutant, which is ~ 5nm shorter than the full-length protein. Grey: molecular dynamics simulation of lipid bilayer^53^ (J) Cartoon depicting a putative dynamic association of VPS13C with the ER membrane due to the flexible linker region of VAP. (K) Proposed model of VPS13C arrangement at ER-endo/lysosome contacts. Surface representation of VPS13C predicted structure reveals a continuous hydrophobic groove (grey) along the protein. For clarity of presentation, some α-helices and disordered loops are not shown.

### Rod-shaped bridges at ER-endo/lysosome contacts

Inspection in cryo-electron tomograms of the space between the ER and endo/lysosomes in VPS13C and VAP-B overexpressing cells revealed abundant rod-shaped densities about 30 nm in length that bridged the two membranes and were perpendicular to them (Fig. 4A, 4A’, 4D). The crowding of these rods in the intermembrane space was consistent with the intensity of the fluorescent signal observed in cells from which these tomograms were obtained. In cells expressing only VPS13C^Halo, where VPS13C accumulates on the entire surface of endo/lysosomes, but does not induce massive ER-endo/lysosome contacts^17^ due to the limited amount of endogenous VAP, no such rods were clearly observed (Fig. 3B, 3F). Most likely, in these cells VPS13C is positioned in different orientations, including an orientation parallel to the endo/lysosomal membrane which would make its detection by cryo-ET very challenging.

To verify that the 30 nm long rods represent VPS13C molecules, we examined cells expressing a truncated form of VPS13C where a portion of the lipid-transfer channel (a.a. 1235-1748) had been removed (Fig. 4B). AlphaFold predictions suggested a length of this construct ~5 nm shorter than full-length VPS13C (Fig. 4I). Accordingly, we found that VPS13C Δ1235-1748 rods between ER and endo/lysosomes were ~24 nm long (Fig. 4E) and that the intermembrane space was ~5nm shorter (i.e. 28.5 nm) than the one mediated by full-length VPS13C (Fig. 4C). Taken together, our comparative cryo-ET analyses of full-length VPS13C and VPS13C Δ1235-1748 support that rod-shaped VPS13C bridges two membranes in situ.

To gain further insight into the characteristics of the rod-like structures, we applied subtomogram averaging and 3D classification analysis. Sub-tomogram averaging without external reference revealed a ~29 nm long and 6 nm wide rod bridging the two membranes (Figs. 4F, S4A and S5). An internal tunnel reflecting the hydrophobic groove could not be seen, but the length and diameter of the rod are in agreement with the structural prediction of VPS13C using AlphaFold, strongly suggesting that the rod is formed by VPS13C. Sub-tomogram averaging of VPS13C Δ1235-1748 revealed ~24 nm long rod (Fig. 4G, 4I), which is also consistent with AlphaFold prediction (Fig. S4). The presence of lipids or the insufficient resolution may obscure such a cavity. Lack of a density that could be assigned to Halo could be explained by the variable orientation of the Halo moiety, which was attached to a flexible loop of VPS13. Likewise, an arc reflecting the VAB domain as shown in low resolution negative staining EM images of purified yeast VPS13^16^ could also not be detected likely because of its variable orientation, as reported in that study. Interestingly, 3D classification analysis (Fig. S4) revealed that in most cases (95% of particles), there was a small gap, ~1-4 nm, between the N-terminal end of the VPS13C rod and the ER membrane (Fig. 4F, 4H, 4J, Movie S4). This gap indicates that the N-terminal region of VAP-anchored VPS13C is not always in direct contact with the ER bilayer. A flexible connection between VAP and VPS13C allowing a small gap accounted primarily by the cytosolic region of VAP is plausible (Fig. 4J): the domain of VAP that binds the FFAT motif (its MSP domain) is at the end of its cytosolic portion which is ~12 nm in length and comprises an unfolded region^36^, while the VPS13C binding site for VAP (its FFAT motif) is localized on an unfolded loop of VPS13C at about 5 nm from the N-terminal end of the rod (Fig. 1B, 1E). We suggest that the direct interaction of VPS13C with the ER bilayer, possibly facilitated by the binding to yet unknown factor(s), is regulated within cells. Alternatively, such unknown factor(s) may be present in limited amount. Altogether, our structural analysis supports a model according to which i) VPS13C functions as a bridge to channel lipids (Fig. 4K), and ii) the association of the N-terminal chorein domain of VPS13C with the bilayer of the ER undergoes regulation.

## Discussion

Our study of VPS13C at ER-endo/lysosome contacts provides the first view of a VPS13-ATG2 family protein in an intact cell and supports the hypothesis that these proteins function as bridges that connect opposite membranes to allow lipid flux between them. The recently published AlphaFold algorithm has allowed us to correlate the predicted full-length structure of VPS13C with the densities obtained by cryo-EM, thus strengthening our conclusions. As VPS13 proteins, which are very large, are expressed at low concentration in cells, in order to visualize with confidence VPS13C-dependent contacts, we had to overexpress VPS13C. Such overexpression may have helped force, by crowding, roughly parallel orientations of VPS13 perpendicular to the two membranes. When expressed at endogenous levels, VPS13 proteins may have more variable orientation and populate contacts with shorter distances between the two membranes if oriented obliquely. VPS13 proteins are very long. Their length may facilitate the formation of bridges when a VPS13 molecule anchored to one membrane explores the space for an appropriate partner membrane. In the case of other membrane tethers, including those that transport lipids by a shuttle mechanism, such property may be achieved by unfolded flexible protein regions present in them.

An unexpected observation made possible by our 3D classification analysis was that, in most cases, the N-terminus of VPS13C, which comprises the chorein domain, appeared to be detached from ER by a 1-4 nm gap, i.e. in a state that does not allow lipid extraction from the bilayer to enter the channel. We suggest that VPS13C may make at least two types of contacts with the ER. One is mediated by VAP, which functions as an adaptor to capture the N-terminal region of VPS13 via an interaction with its FFAT motif, but does not force a direct contact of VPS13C with the ER bilayer because of the long and flexible linker region in VAP. The other one, supported by the occurrence of a subpopulation of VPS13C rods in direct contact with the ER membrane, could be a regulated, possibly low affinity, interaction of the highly conserved N-terminal chorein domain of VPS13C with some component (protein or lipid) of the ER membrane. Binding of VPS13C to VAP may facilitate the occurrence of this other interaction. We note that the chorein domain is the most conserved region in VPS13-ATG2 family proteins, leading to the speculation that such a motif is critically important for membrane binding and lipid extraction. Yet, ATG2 and SHIP164, which have very similar chorein motifs^23, 29^, but lack FFAT motifs for VAP binding, do not have an obvious localization throughout the ER even when overexpressed^24, 37^. Likewise, VPS13A and VPS13C lose their ER association if their FFAT motif is deleted^17^. Thus, VAP-independent ER interactions of VPS13C must be of low affinity, regulated and transient. It was proposed that VMP1 and TMEM41, which have scramblase activity, represent binding sites in the ER for ATG2^38^. Along with this possibility, VMP21 was shown to function upstream of VPS13D in Drosophila^39^. However, so far, we have not detected an interaction of VPS13 with these proteins.

A dynamic association of VPS13 with the ER bilayer may be a mechanism that contributes to a precise regulation of bulk lipid transfer to other membranes. Further elucidation of these mechanisms may not only advance fundamental knowledge about cell biology but also provide insight into how mutations of these proteins lead to neurodegenerative and neurodevelopmental disorders.

## Methods

### DNA plasmids

A plasmid containing codon-optimized cDNA encoding full-length human VPS13C, with a Halo protein after amino acid residue 1914 (VPS13C^Halo), was generated by and purchased from GenScript Biotech. GFP-VAPB and PDZD8-EGFP were previously generated in our laboratory^34, 40^. A deletion mutant of VPS13C lacking amino acid 1235-1748 was generated as follows. First a C-terminal fragment of VPS13C^halo_1749-3753_, including a Halo protein was generated by PCR amplification from VPS13C^halo plasmid and ligated into pEGFP-C1 by InFusion Cloning using the EcoRI and KpnI sites. Then VPS13C^Halo (Δ1235-1748) construct was generated by PCR amplification from VPS13C^Halo plasmid and ligated into VPS13C^halo_1749-3753_ plasmid by InFusion Cloning using the NheI and EcoRI sites.

### Cell culture and transfection

HeLa cells were cultured in DMEM (Thermo Fisher Scientific) supplemented with 10% FBS (Thermo Fisher Scientific) and maintained at 37°C, 5% CO2 in a humidified incubator. For transfection, cells were seeded on 35mm Petri dishes at a concentration of 50,000 cells per dish and transfected after 24h using FuGene HD (Promega).

### Cell culture on EM grids

Preparation of EM grids for cell seeding was performed as previously described, with modifications^41^. Four to six Quantifoil R2/4 Au-grids with carbon sides facing up were placed onto the center of a 35 mm glass-bottomed dish (MatTek, 0.15 mm in thickness and 2 mm in diameter). The carbon side of the grids were glow discharged for 45 s at 15 mA in a plasma cleaner (PELCO easiGlow, Ted Pella, Redding, CA, USA). The dish with grids was then sterilized in 5ml 70% ethanol for 10 min under UV, washed with distilled water for six times and incubated with 1ml 0.1 mg/ mL poly-D-lysine (Thermo Fisher Scientific) overnight in a 37 °C incubator. The grids were subsequently washed with distilled water for six times and incubated with DMEM complete media overnight.

Right before seeding cells, each grid was placed onto the center of a separate glass-bottomed dish (MatTek). Transfected cells were trypsinized and 1500 cells were seeded on each grid. The grid was incubated at 37°C, 5% CO2 for 10 mins and then 2ml DMEM complete media was slowly added to each dish.

### Live-cell fluorescence microscopy

Cells transfected with VPS13C^halo and GFP-VAP were seeded on EM grids on glass-bottomed MatTek dishes one day prior to fluorescence imaging. Halo tag ligands JF549 were added to each dish at a final concentration of 200 nM. Cells were incubated with the dye for 1 h, rinsed for three times, and then incubated in DMEM complete media for 1 h before imaging. Spinning-disk confocal imaging of the cells was performed using an Andor Dragonfly spinningdisk confocal microscope equipped with a plan apochromat objective (63x, 1.4 NA, oil) and a Zyla scientific CMOS camera at 37°C and 5% CO2. The grid was placed in an orientation with the “1” label at the top. Cells with bright VPS13C and VAP fluorescence were imaged and their location relative to the center of the grid was recorded.

### Conventional Correlative Light and Electron Microscopy

For transmission electron microscopy CLEM, Cos7 cells were plated on 35 mm MatTek dish (P35G-1.5-14-CGRD) and transfected with VPS13C^Halo, EGFP-VAP-B, PDZD8-EGFP, and mCherry-Rab7 constructs. Cells were pre-fixed in 4% PFA+0.25% glutaraldehyde in Live Cell Imaging Buffer (Life Technologies) then washed before fluorescence light microscopy imaging. Regions of interest were selected and their coordinates on the dish were identified using phase contrast. Cells were further fixed with 2.5% glutaraldehyde in 0.1 M sodium cacodylate buffer, postfixed in 2% OsO4 and 1.5% K4Fe(CN)6 (Sigma-Aldrich) in 0.1 M sodium cacodylate buffer, en bloc stained with 2% aqueous uranyl acetate, dehydrated, and embedded in Embed 812. Cells of interest were relocated based on the pre-recorded coordinates. Ultrathin sections (50-60 nm) were observed in a Talos L 120C TEM microscope at 80 kV, images were taken with Velox software and a 4k × 4K Ceta CMOS Camera (Thermo Fisher Scientific).

For correlative light microscopy and Focused Ion Beam Scanning Electron Microscopy (FIB-SEM), VPS13C^Halo and PDZD8-EGFP were co-expressed in Cos7 cells and processed as above with the exception that except fluorescence light microscopy imaging was carried out in live mode without the pre-fixation with PFA. Epon blocks were glued onto the scanning electron microscope mounting aluminum stub. Next, platinum en bloc coating (20-25 nm thick) of the sample surface was carried out with the sputter coater (Ted Pella, Inc., Redding, California). Sample were FIB-SEM imaged in a Crossbeam 550 FIBSEM workstation operating under SmartSEM (Carl Zeiss Microscopy GmbH, Oberkochen, Germany) and Atlas 5 engine (Fibics incorporated, Ottawa, Canada). The imaging resolution was set at 7 nm/pixel in the X, Y axis with milling being performed at 7 nm/step along the Z axis to achieve an isotropic resolution of 7nm/voxel. Images were aligned and exported with Atlas 5 (Fibics incorporated, Ottawa, Canada), further processed and analyzed with DragonFly Pro software (Object Research Systems (ORS) Inc., Montreal, Canada). Except when noted all reagents were from EMS (Electron Microscopy Sciences), Hatfield, PA.

### Plunge freezing

Dishes with grids were taken out of the incubator and the media of each dish was replaced with 1ml live-cell imaging solution (Life Technologies) to maintain physiological pH. Plunge freezing was performed with Vitrobot Mk IV (Thermo, Waltham, MA). The Vitrobot was set to 37°C, 100% humidity. The grid seeded with cells was held with a Vitrobot tweezer, immersed in a livecell imaging solution with 3-5% glycerol for 1 min and then mounted onto the Vitrobot and blotted once in the Vitrobot chamber (blot force: 10, blot time: 10s). To ensure that cells on the grids were not too dry, the side of the chamber facing the carbon side of the grid (where cells are located) was covered with two layers of parafilm while the side facing the gold side of the grid was covered with two layers of blotting paper (Whatman). The grid was then plunged into a liquid ethane-propane mixture (4:6 ratio).

### Cryo-focused ion beam milling

Micromachining of frozen hydrated cells was performed in FEI Aquilos1 and Aquilos2 FIB/SEM systems, operated at a temperature below −180°C. Grids were imaged by scanning EM imaging (5 kV, 21 pA) to check quality of the sample. Samples were sputter coated with a platinum metal layer for 15s. Then Gas-Injection-System (GIS) deposition of organo-platinum was applied for 10-15s. Cryo-lamellae were milled in several steps. A 9-μm-wide and 3-μm-thick cryo-lamellae was first generated with the ion beam set to 30 kV and ~500 pA. The ion-beam current was then decreased gradually until the cryo-lamellae reached a nominal thickness of 500 nm (ion beam set to 50 pA). After the thinning of all cryo-lamellae, each cryo-lamella was polished to a nominal thickness of ~100-150 nm (ion beam set to 10 or 30 pA). An SEM image of each cryolamella was taken at 5 kV, 30 pA, 1μs dwell time. The polishing step was completed within an hour to minimize contamination. A thin platinum metal layer was then applied (4 to 5 second sputter coating) to improve the electrical conductivity of the cryo-lamellae surface if phase-plates were used during the following TEM imaging.

### Cryo-electron tomography

Tomographic tilt series were recorded in a Titan Krios TEM equipped with a field emission gun operated at 300 kV, a GIF Quantum LS post-column energy filter (Gatan) and a K3 Summit direct electron detector (Gatan). Tilt series were acquired at a target defocus of −6 μm and a pixel size of 3.3Å using the SerialEM software in low-dose mode. The dose-symmetric tomography acquisition scheme was applied and modified according to the pre-tilt of the lamella. The K3 detector was operated in counting mode and 10 sequentially acquired 0.1 sec frames were combined. The tilt range was typically between ±50°and ±60°with increments of 3°and the total electron dose was around 80 e/Å^2^. Imaging details are shown in Table S1.

### Tomogram reconstruction

The frames of each tilt series micrograph were aligned using MotionCor2^42^. For defocus-contrast micrographs, contrast transfer function (CTF) correction was performed in Gctf ^43^. A script written by Yan Rui from the HHMI Janelia CryoEM Facility was used to reorder and combine tilt series micrographs. Alignment of the tilt series and tomographic reconstructions were performed using Etomo, which is part of the IMOD package^44^. Fiducial tracking was performed with 5-10 nm-sized gold-bead-like deposits on the lamella. Tomograms were reconstructed using the weighted back-projection (WBP) and Simultaneous Iterative Reconstruction Technique (SIRT). Tomograms were 4x-binned for subsequent analysis. For figure presentation of defocus-contrast tomograms, either a Nonlinear Anisotropic Diffusion filter^44^ or a deconvolution filter (https://github.com/dtegunov/tom_deconv^45^) was applied to enhance contrast. Segmentation of ER, endo/lysosome membranes, microtubules was done in IMOD^44^. Segmentation of ribosomes was done in EMAN2^46^. More details of image analysis and the datasets used in this paper are shown in Table S1 and Table S2, respectively.

### Subtomogram averaging and 3D classification

Rod-shaped VPS13C-like densities were manually selected (Full-length VPS13C: 576 in total; VPS13C Δ1235-1748: 110 in total) from 4x binned tomograms using IMOD, with the first point on the tip of the VPS13C rod close to ER and the second point on the tip at the opposite side, next to the endo/lysosome. Subtomograms of the selected densities were then extracted from 4x or 2x binned tomograms and the i3 software package was used for 3D multi-reference alignment and classification, which were computed in Fourier space^47, 48^. To estimate the resolution of the averaged density map, Fourier shell correlation (FSC) coefficients were calculated with the i3 software package and plotted with Excel. The remapping of rods to the original tomogram was done using the scripts i3_to_RELION.py (https://github.com/scai20/i3) and ot_remap.py (https://github.com/anaphaze/ot-tools) and visualized with UCSF Chimera ^49^. Rigid-body fitting of AlphaFold predicted structure into the averaged density map was done in UCSF ChimeraX^50^.

### Measuring distances between membranes

Segmentation of ER and endo/lysosome membranes in close contact with each other was done in IMOD^44^. The coordinates were subsequently extracted. The nearest-neighbor distance between multiple points on the ER membrane and multiple points on the endo/lysosome membrane within the segmented area was calculated and the distribution was plotted with a custom python script (https://github.com/scai20/Intermembrane-space). A violin plot was generated with a Python data visualization library Seaborn^51^.

### AlphaFold structure prediction

Structural predictions of three segments of VPS13C (a.a. 1-1860, 1201-2340, 1801-3753) were generated with AlphaFold v2.0^29^ on the Yale Farnam high performance computer cluster. The three fragments were aligned and combined in UCSF ChimeraX^50^. Alphafold-Multimer^30^ was used to predict the interaction between VPS13C N-terminus and VAP.

### Data Sharing

Subtomogram-average density maps showing a full-length VPS13C rod and a VPS13C Δ1235-1748 rod bridging the two adjacent membranes have been deposited at EMDB as entry EMD-26247. The raw cryo-ET tilt series and reconstructed tomograms presented in Figures 3, 4 and S2 have been deposited in EMPIAR ^52^ under entry EMPIAR-10962.

## Acknowledgments

We thank Marianna Leonzino, Xia Li, Lu Gan (National University of Singapore) for advice and discussion. We also acknowledge the support of the cryo-EM facilities of the Yale West Campus, of the Yale Center for Cell Imaging of the Yale Medical School and of the HHMI Janelia Research Campus. This work was supported in part by grants from the NIH (DA018343, NS36251 to P.D.C. and AI152421 to J.L.), and by the Kavli Institute for Neuroscience, the Parkinson’s Foundation, the Chan Zuckerberg Initiative DAF, an advised fund of Silicon Valley Community Foundation and the Aligning Science Across Parkinson’s grant ASAP-000580 through the Michael J. Fox Foundation for Parkinson’s Research (MJFF) to P.D.C. For the purpose of open access, the authors have applied a CC BY public copyright license to all Author Accepted Manuscripts arising from this submission. S.C was supported by a fellowship from the Parkinson’s Foundation. P. De Camilli serves on the scientific advisory board of Casma Therapeutics.

---

**Figure S1.**
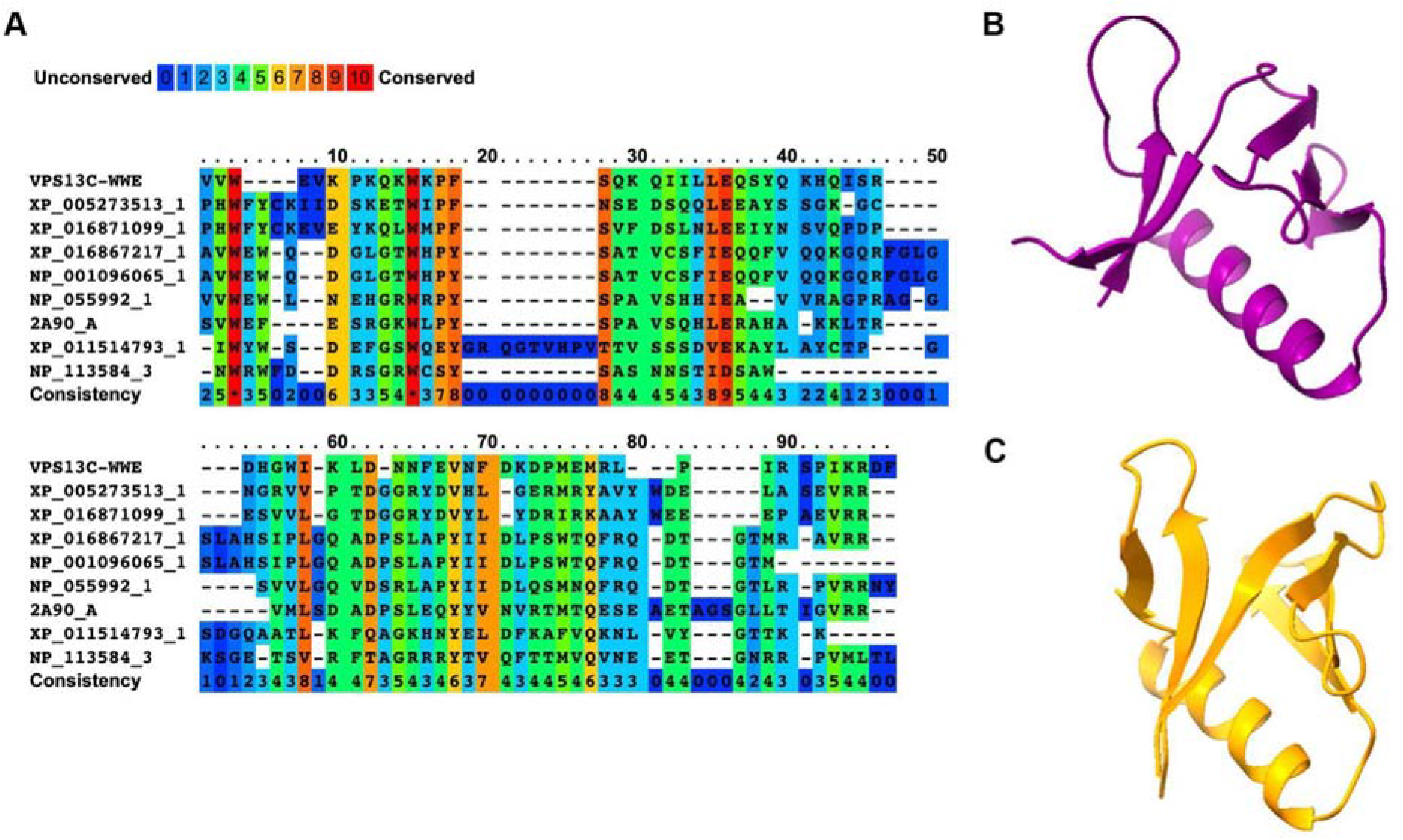
A fragment of VPS13C (a.a 3117-3183) aligns with WWE domains. (A) Sequence alignment of a VPS13C fragment (a.a 3117-3183) with WWE domains of multiple proteins by PRALINE multiple sequence alignment^54^. (B) Structural prediction of VPS13C3117-3183 by AlphaFold v2.0 (C) Structure of WWE domain of human HUWE1 (PDB 6MIW).

**Figure S2.**
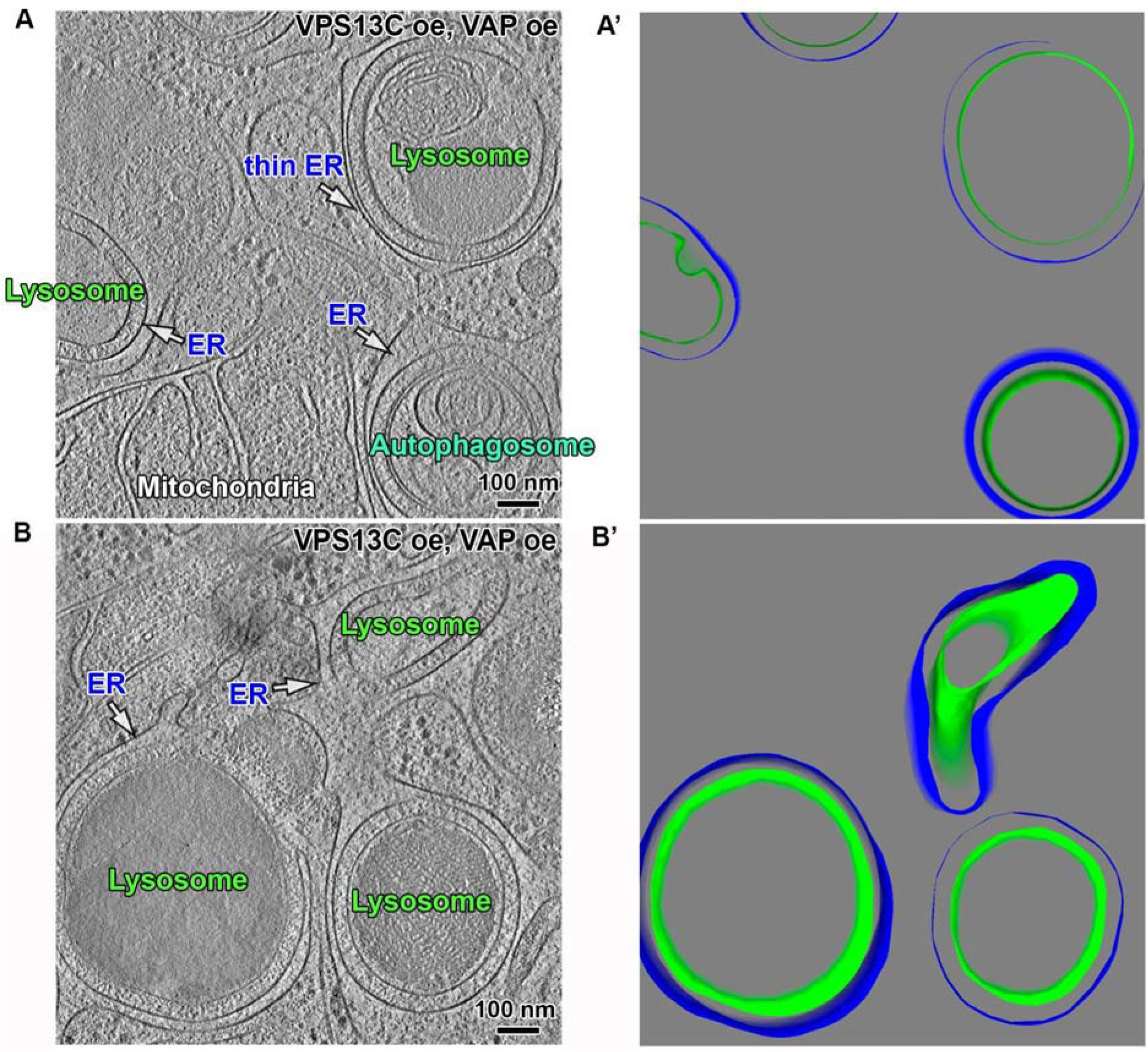
Examples of VPS13C mediated contact sites. (A and B) Cryotomographic slices (1-nm thick) showing VPS13C-mediated membrane contacts in HeLa cells overexpressing VPS13C^Halo and GFP-VAP-B. (A’ and B’) 3D segmentation views of panels A and B, respectively, showing endo/lysosome membranes in green, an autophagosome membrane in cyan and ER membranes in blue. intermembranes spaces are partially hidden by the 3D representation.

**Figure S3.**
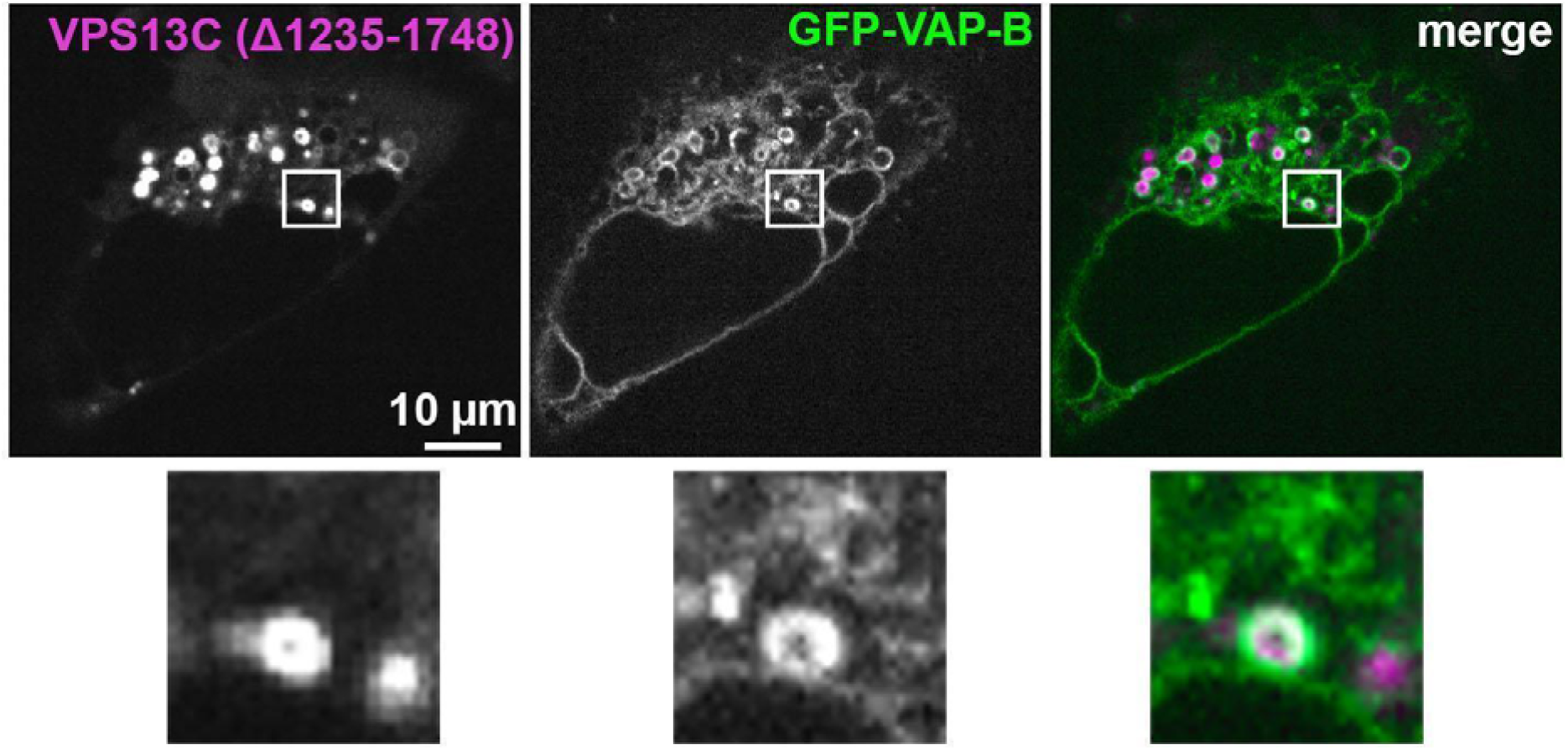
The VPS13C truncation mutant (Δ1235-1748) has the same subcellular localization as full-length VPS13C. Confocal microscopy z-slice images of a HeLa cell expressing VPS13C^Halo (Δ1235-1748) and GFP-VAP-B in the split and merged channels. The region enclosed by a box is shown at five fold enlargement at the bottom.

**Figure S4.**
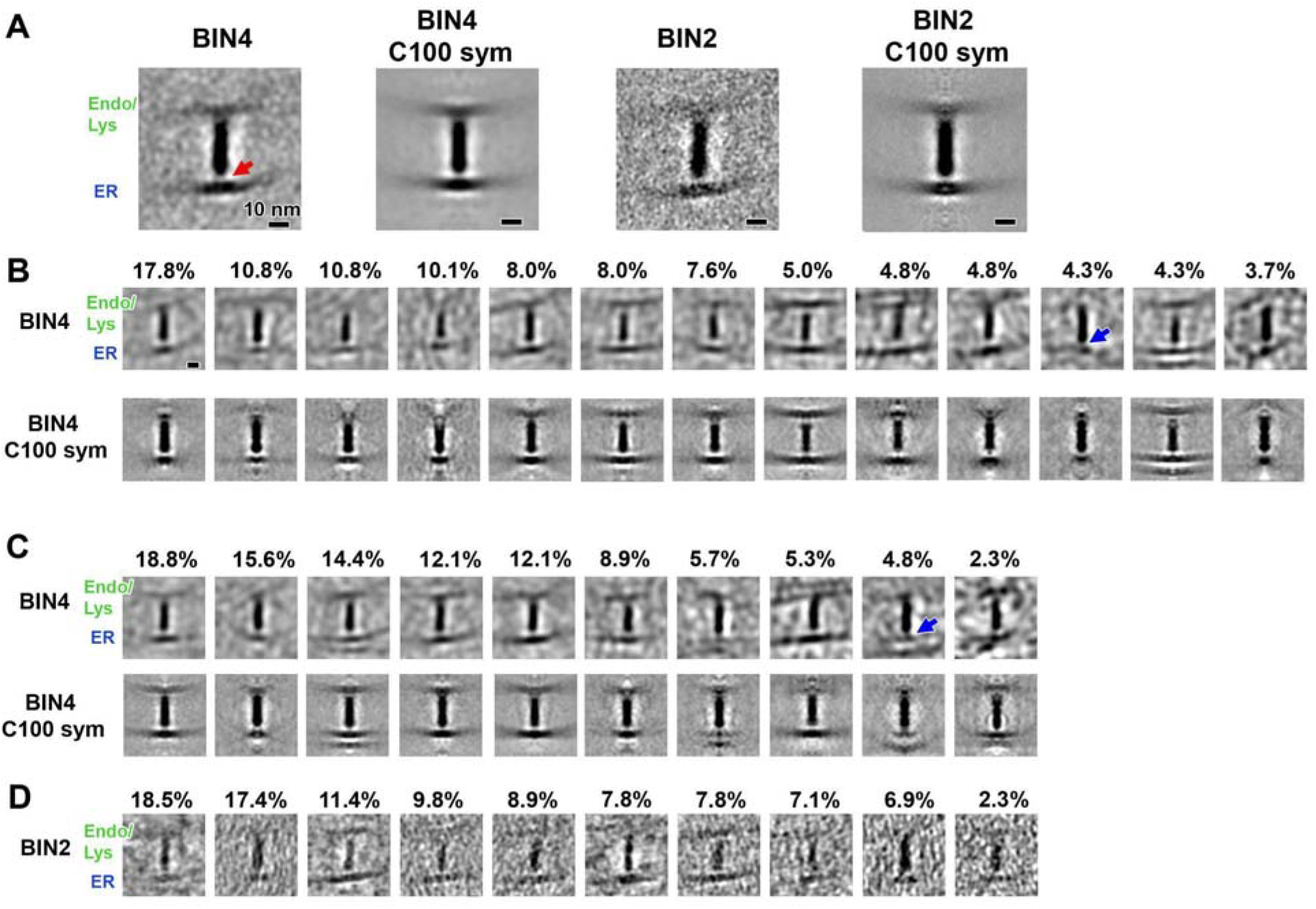
Subtomogram averaging and 3D classification. (A) Subtomogram averages with subtomograms binned by 4 and binned by 2, respectively. C100 sym: averages symmetried with C100 to enhance signal to noise ratio. Red arrow: a gap between the N-terminal end of the VPS13C rod and the ER membrane (B) Class averages (3D) of subtomograms binned by 4. Blue arrow: a class showing no obvious gap between the N-terminal end of the VPS13C rod and the ER membrane (C) Class averages (3D) of subtomograms binned by 4. A mask around ER side of the rod was applied. Blue arrow: a class showing no obvious gap between the N-terminal end of the VPS13C rod and the ER membrane (D) Class averages (3D) of subtomograms binned by 2.

**Figure S5.**
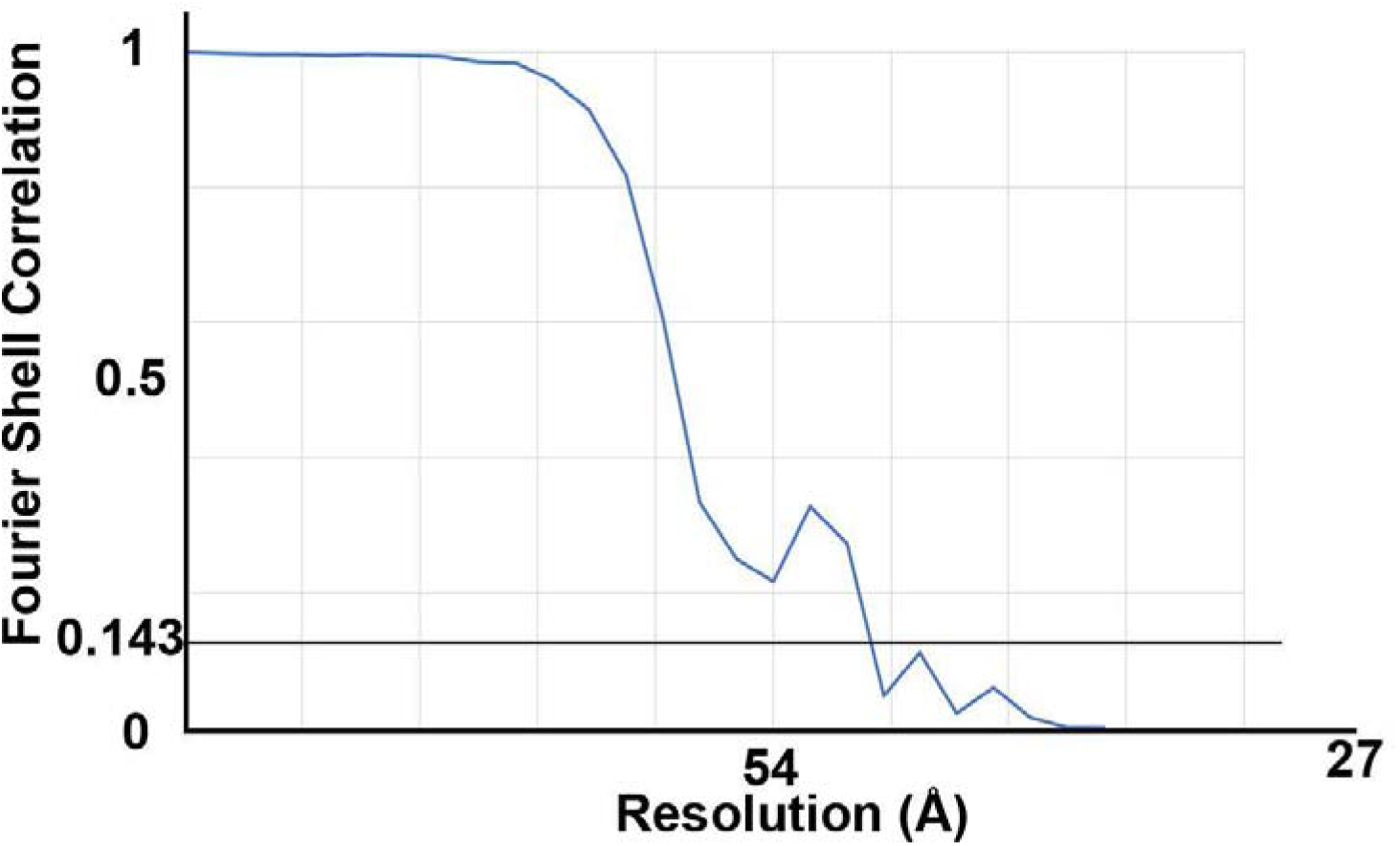
Fourier Shell Correlation of the structure shown in Figure 4F. The resolution of the averaged density map in Figure 4F is ~ 47 Å, based on the Fourier Shell Correlation = 0.143 criterion.

**Movie S1. Predicted structure of full-length VPS13C bound to the MSP domain of VAP.**

The color scheme is the same as the one shown in Fig 1A.

**Movie S2. Predicted structure of full-length VPS13C showing a continuous hydrophobic groove**

Surface representations of the structure of VPS13C predicted from AlphaFold showing carbon atoms in grey (hydrophobic surfaces), oxygens in red (negative charges), nitrogens in blue (positive charges) and sulfur in yellow. For clarity of presentation, some α-helices and disordered loops are not shown.

**Movie S3. FIB-SEM image stack of endo/lysosomes from Cos-7 cells overexpressing VPS13C^halo and PDZD8-EGFP**

FIB-SEM image stack of the structure shown in Fig. 2E’. VPS13C and PDZD8 segregate to distinct domains within the contact area between ER (thin ER) and the endo/lysosome.

**Movie S4. Three-dimensional view of VPS13C averaged density map**

Three-dimensional view of the VPS13C rod bridging two membranes. The density corresponding to VPS13C is shown in light orange, fitted with the full-length VPS13C predicted structure from AlphaFold v2.0. The color scheme of predicted structure is the same as that of Fig. 1A. The densities corresponding to ER and endo/lysosome membrane are colored in blue and green, respectively.

**Table S1.** Cryo-ET and image-analysis details.

|  |  |
| --- | --- |
| EM grid | Quantifoil R 2/4, LF, 200 Mesh, Gold |
| Microscope | Titan Krios |
| Energy | 300 kV |
| Camera: recording mode | Gatan K3 Summit: counting |
| Subframes / tilt | 10 |
| Energy filter / zero-loss slit width | GIF energy filter/ 20 eV |
| Tomography software | SerialEM |
| Calibrated pixel size, unbinned | 3.3 Å |
| Defocus (nominal, defocus contrast) | -6 µm |
| Defocus (nominal, Volta contrast) | -1 µm |
| Cumulative dose | 80 e<sup>-</sup> / Å<sup>2</sup> |
| Dose fractionation | constant |
| Tilt range | ±50-60°, dose-symmetric |
| Tilt increment | 3° |
| Movie alignment | MotionCor2 |
| Tomogram processing & visualization | IMOD 4.11 |
| Segmentation | IMOD 4.11, EMAN 2.91 |
| Subtomogram classification & averaging | I3 0.9.9.3, RELION 3.0.4 |
| Density map visualization | UCSF ChimeraX, UCSF Chimera |
| Figure/movie creation and editing | Adobe Photoshop, Illustrator |

**Table S2.**
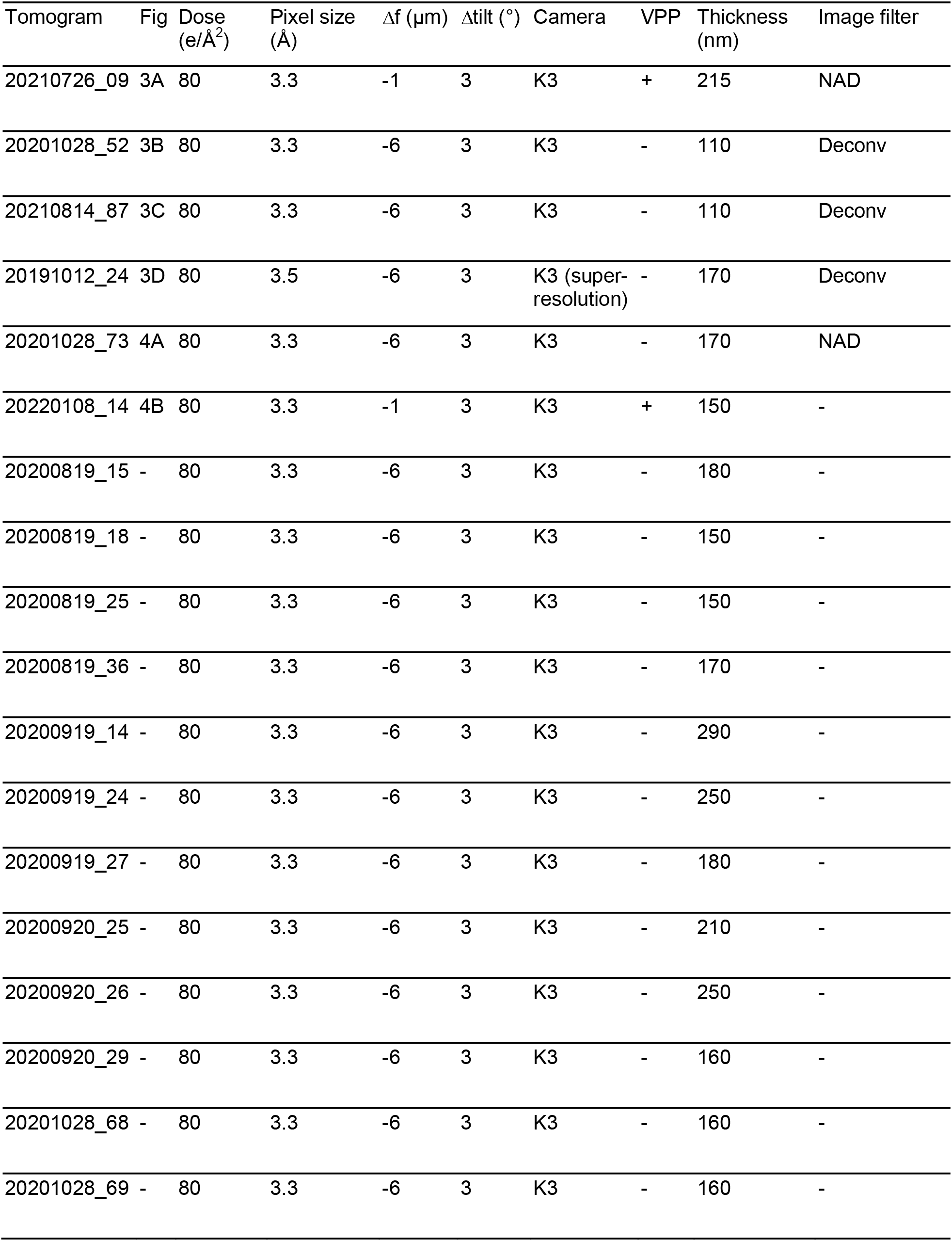
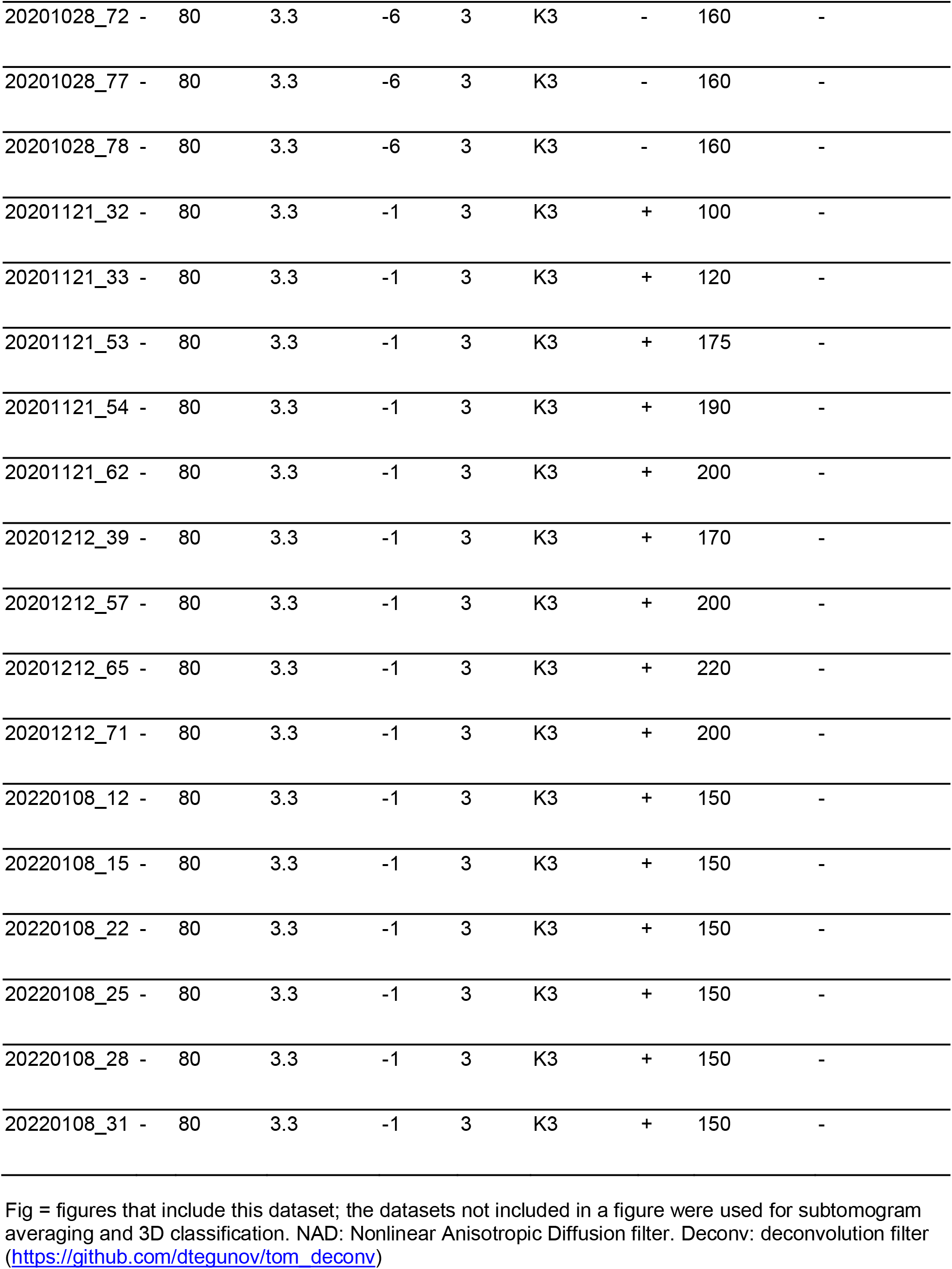
Cryotomogram details.

## Notes

### Competing Interest Statement

The authors have declared no competing interest.

